# A Novel Technique for Fluorescence Lifetime Tomography

**DOI:** 10.1101/2024.09.19.613888

**Authors:** Navid Ibtehaj Nizam, Vikas Pandey, Ismail Erbas, Niels Bracher, Jason T. Smith, Xavier Intes

## Abstract

Fluorescence lifetime imaging has emerged as a powerful tool for quantitatively assessing the molecular environment of live tissues *in vivo*. While fluorescence lifetime microscopy (FLIM) is a mature field, achieving effective 3D imaging in deep tissues has remained a significant challenge due to high scattering. In this study, we present a deep neural network-based approach, referred to as AUTO-FLI, which enables both 3D intensity and quantitative lifetime reconstructions at centimeters depth. This Deep Learning (DL)-based method incorporates an *in silico* framework to accurately generate fluorescence lifetime data for training and validation. The performance of this novel DL model is further validated with experimental data acquired on an anatomically accurate mouse-mimicking phantom. The results demonstrate that AUTO-FLI can provide precise 3D quantitative estimates of both intensity and lifetime distributions in highly scattering media. This method holds great promise for fluorescence lifetime-based molecular imaging at both the mesoscopic and macroscopic scales, with potential applications for pre-clinical translational research and fluorescence-guided surgery.

## I. Introduction

IN recent years, molecular imaging has become a vital tool for investigating biological processes *in vivo* in both pre-clinical and clinical settings. Due to its high sensitivity, availability of a wide range of fluorophores, and multiplexing capabilities, fluorescence optical imaging has emerged as the most prominent molecular imaging technique in the last decade [1], [2]. Although intensity (brightness) remains the primary contrast mechanism in fluorescence optical imaging, fluorescence lifetime is increasingly leveraged for its unique advantages. Fluorescence Lifetime Imaging (FLI) can quantitatively sense various intracellular parameters, including metabolic state, viscosity, temperature, and pH [3]–[5]. Additionally, fluorescence lifetime allows for robust quantification of Förster Resonance Energy Transfer (FRET), enabling nanoscale assays *in vivo* and the study of drug-target engagement [6]–[8].

FLI has grown steadily over the past three decades, with increased adoption due to user-friendly FLI Microscopes (FLIM) [9], [10]. Simultaneously, FLI has expanded to translational applications, from mesoscopic (mFLI) [11] to macroscopic regimes (MFLI) [8], [12], [13]. These larger-scale implementations are more challenging than microscopic ones because they require Near-Infrared (NIR) fluorophores for deeper tissue penetration. Red-shifted fluorophores typically have shorter lifetimes (nanosecond or sub-nanosecond) compared to a few nanoseconds in the visible range, and detectors have low quantum efficiency (a few percent) [14]. Furthermore, 3D lifetime imaging in deep tissues is hindered by light scattering. Achieving 3D quantitative maps in these regimes requires solving a diffuse optical inverse problem, which considers the spatial and temporal characteristics of illumination light, its propagation, energy transformation, temporal delays generating fluorescence photons, and the propagation of these photons to the detection surface. This problem is computationally challenging and extremely difficult to solve using classical methods, limiting mFLI and MFLI mainly to 2D imaging applications.

In principle, 3D Fluorescence Lifetime Tomography (FLT) is a “double ill-posed problem” because FLT involves reconstructing a 3D volume from a scattering medium while simultaneously determining the lifetime for each voxel. Two main approaches have been proposed to simplify the complexity of this inverse problem: estimating the lifetime from 2D measurements before casting the inverse problem using this prior information or linearizing the lifetime exponential model to stabilize the inverse problem. Pertaining to the first method, Kumar et al. [15] proposed an asymptotic approach in which 2D fluorescence data are preprocessed to obtain the relative intensities associated with individual fluorophores, and then the inverse problem is cast independently for each fluorophore using a quasi-Continuous Wave (CW) formulation. However, this asymptotic assumption is valid only for long lifetime values (*τ >* 0.5 ns) and does not consider the time-of-flight delays associated with scattering and tissue depth. This approach was also combined with structural priors obtained by Computational Tomography (CT) to constrain the inverse problem further [16]. The integration of CT data increases the complexity and cost of the imaging process while also enforcing congruity between the two modalities, which is seldom attained biologically. Furthermore, Zhang et al. [17] used a mathematical technique called the fused LASSO method, which involves reconstructing the fluorescence yield map and object geometry as prior information, helping to mitigate the ill conditions associated with the nonlinear physical model of FLT. However, this linear approximation is computationally complex and highly dependent on setting regularization parameters. Another approach proposed by our group involves estimating lifetimes from 2D measurements and incorporating these values as priors in the 3D inverse problem, utilizing the rich information of time-resolved datasets [18]. Although this approach is technically more precise and can offer improved resolution thanks to the incorporation of early photons [19], [20], it leads to increased computational burden (size of the inverse problem to solve). Hence, common limitations of current methodologies reside in their time-consuming computations, simplifications that are valid for limited cases, and heavily reliant on hyperparameter optimization. Following recent trends, Deep Learning (DL) methodologies are expected to provide efficient frameworks to overcome these challenges [21], [22].

With the advent of high-powered Graphical Processing Units (GPUs), the use of Deep Neural Networks (DNNs) in Fluorescence Molecular Tomography (for 3D intensity reconstructions) has flourished in recent years. Long et al. proposed one of the first networks to achieve FMT reconstructions in the mesoscopic domain, employing a CNN. Other strategies by Zhang et al. [24] utilized CNNs with skip connections to achieve high-resolution FMT reconstructions. Previously, we introduced a novel architecture called ModAM [25], which combined a *k*-space illumination basis and a unique *in silico* data generation workflow for 3D FMT reconstructions [26]. However, despite the proliferation of DNNs specializing in FMT intensity reconstructions, no solutions have been proposed for achieving 3D lifetime reconstructions with high fidelity. The lack of progress is mainly due to the high difficulty in designing a network that accurately learns the lifetime from the raw fluorescence decays while simultaneously reconstructing a 3D inclusion. Furthermore, the reliance on experimental data for training is a massive bottleneck, as collecting a large experimental dataset for different physical and Optical Properties (OPs) involved with fluorescence lifetime estimation is highly time-consuming and serves as a deterrent to designing a network that can achieve 3D FLT reconstructions.

Here, we propose AUTO-FLI, a CNN that performs end-to-end 3D FMT reconstructions for relative quantum yields and lifetimes. To our knowledge, this is the first DL-based technique to achieve full 3D volumetric reconstruction of lifetime from raw 2D fluorescence decays. The two-stage workflow involves 3D quantum yield reconstruction using the ModAM network, which constrains the 3D lifetime output from a modified FLI-NET. Using a mono-exponential lifetime model (for simplicity and a first analysis), we deploy a *k*-space illumination basis [26] and generate training data with Monte-Carlo eXtreme (MCX), a MATLAB-based simulator [27]. This approach ensures accurate modeling of fluorescence imaging physics and mitigates the need for extensive experimental data. AUTO-FLI is validated on *in silico* and experimental data from a tissue-mimicking phantom. Additionally, to demonstrate the algorithm’s suitability for pre-clinical applications, we test the performance of the proposed technique on a biologically accurate mouse phantom. The *in silico* results are evaluated for Mean Squared Error (MSE) and Volume Error (VE), while the experimental results are benchmarked against a state-of-the-art 2D fluorescence lifetime estimation software, AlliGator [28].

## II. Methods

### A. Data Generation Workflow

The data generation workflow is illustrated in Fig. 1. To generate the training dataset, we first design *in silico* phantoms in MCX (Fig. 1B). These phantoms contain one, two, or three fluorescent inclusions, which are randomly picked from our previously proposed EEMINST dataset (Fig. 1A) [26]. This dataset is designed to have high spatial heterogeneity, enabling our network to learn to reconstruct complex structures (such as vascular beds or spiculated cancers). Moreover, the fluorescent embedding(s) (with thicknesses varying from (2–4) mm) are placed at different depths (1–10) mm within the phantom. In the case of multiple fluorescent inclusions, the relative concentration of the species ranges from 1 : 1 to 1 : 4. Additionally, the background is given OPs relevant to soft tissues with an absorption coefficient (*μ*_*a*_) of 0.004 mm^*−*1^ and a reduced scattering coefficient 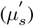 of 1 mm^*−*1^. Next, we use widefield, *k*-space illumination patterns (the same patterns used in Nizam et al. [26], Fig. 1D) acquired experimentally from our gated-Intensified Charged-Couple Device (ICCD) system (discussed in the experimental setup section) to illuminate the phantom (Fig. 1C). Through MC simulation, the flux at the position of the embeddings (*ϕ*) is gathered and then multiplied with the relative concentration matrix (1 : 2 as an example in Fig. 1E) to obtain *ϕ*_*c*_ (shown for a single *k*-space pattern in Fig. 1F). In the next step *ϕ*_*c*_ is convolved with an exponential decay profile corresponding to a mono-exponential lifetime ranging from (0.3–2.5) ns (an example lifetime map is given in Fig. 1G), which results in a time-varying profile, *ϕ*_*t*_ (shown for a single *k*-space pattern in Fig. 1H).

**Fig. 1.**
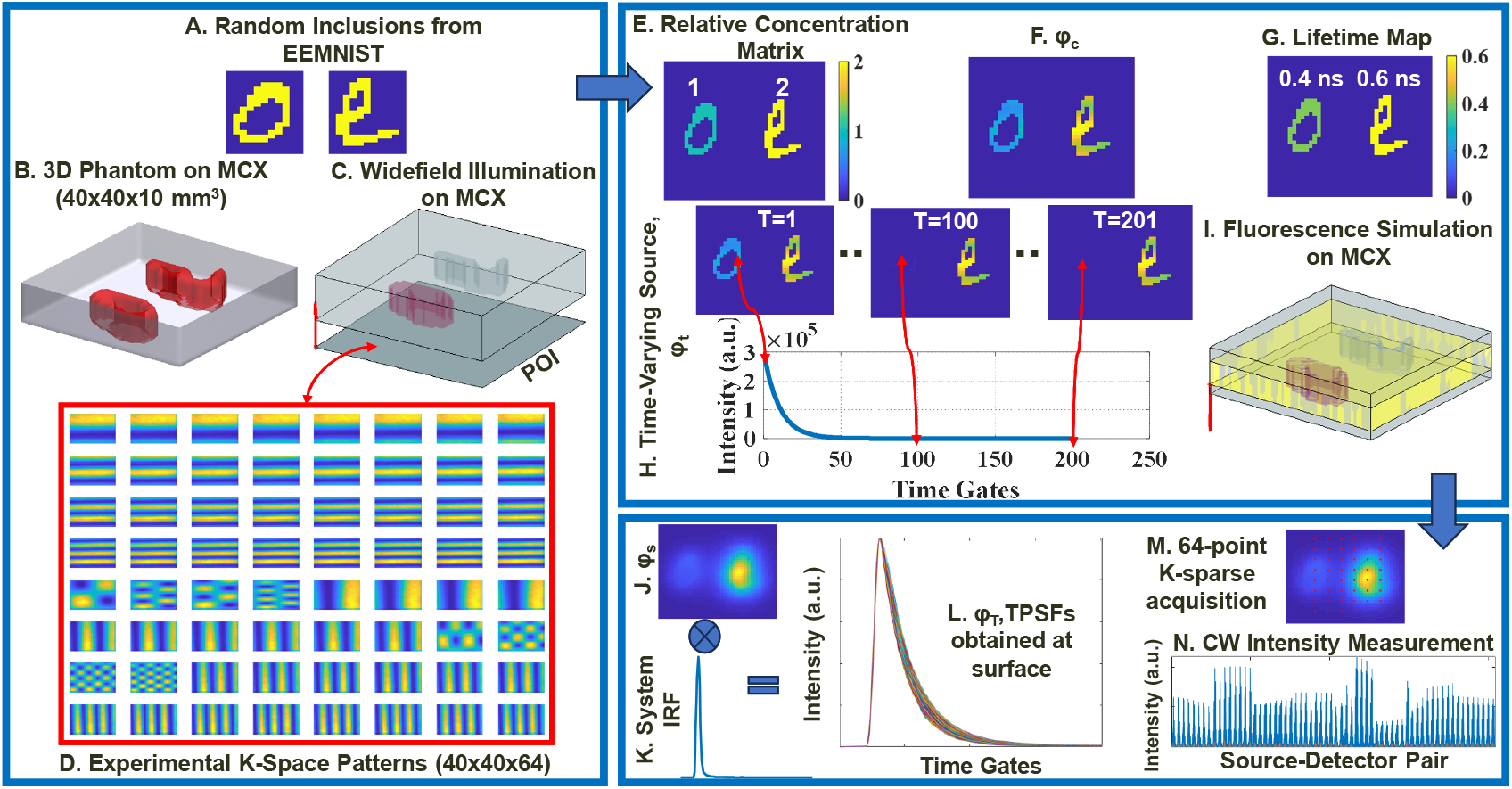
The complete data generation workflow with an example *in silico* phantom in B containing two fluorescent inclusions randomly picked from the EEMNIST dataset shown in A. This phantom is illuminated using K-space illumination patterns (on the Plane of Illumination (POI) marked in C) acquired experimentally, as shown in D. The widefield illumination function in MCX shown in C allows the flux at the position of the embeddings to be collected, which is correlated with the concentration matrix in E to obtain ***ϕc*** in F. F is convolved with a mono-exponential lifetime profile associated with the lifetime map in G to obtain H as a time-varying source. H is used as a propagator from inside the medium on MCX, as shown in I, to obtain J, which is convolved with the system IRF in K to obtain the TPSFs in L and through 64-point K-space acquisition on K, as illustrated by the red dots in M, the CW measurement vector is obtained in N.

This *ϕ*_*t*_ (obtained for each *k*-space pattern) is now used as a propagator from inside the medium (in the opposite direction to simulate fluorescence) and again through MC simulation (Fig. 1I), the flux at the surface of the phantom, *ϕ*_*s*_ (shown for a single *k*-space pattern in Fig. 1J), is collected. Finally, *ϕ*_*s*_ is convolved with the Instrument Response Function (IRF) (obtained experimentally using the gated ICCD system from a thin sheet of paper placed at the same height as the surface of the experimental phantom and shown for a single pixel in Fig.1K), to obtain the Temporal Point Spread Functions (TPSFs), *ϕ*_*T*_ (shown normalized for a single *k*-space pattern in Fig. 1L). Only the falling edge of the normalized TPSFs (corresponding to the decay) is preserved for training the network. Also, *ϕ*_*T*_ is summed in time, obtaining a CW map for each pattern, and using *k*-sparse acquisition (with the help of 64 virtual point detectors, as shown in Fig. 1M) [26]), we obtain a CW measurement vector for the phantom (shown in Fig. 1N). This CW measurement vector is used to train the ModAM branch of the network for 3D intensity reconstruction. The physical dimensions shown in Fig.1 are those used in the data generator to train the network to reconstruct the experimental phantom. These dimensions are suitably adjusted to train the data generator for the mouse phantom.

### B. CNN Architecture and Training

Our proposed AUTO-FLI network comprises two main branches (Fig. 2): the ModAM branch to reconstruct the 3D intensity and the modified FLI-NET branch for reconstructing the 3D lifetime. The final lifetime output is constrained by the 3D intensity mask so that only regions with fluorescent activity are considered in the 3D lifetime map. Both branches combine 2D and 3D convolutional layers, each followed by batch normalization and a ReLU activation, except for the final layer in each branch. The rationale for the various layers in the neural network architecture has been detailed in previous work [26], [29]. For clarity, we will provide a brief overview summarizing their roles in our setting and provide the mathematical framework for the proposed AUTO-FLI network.

**Fig. 2.**
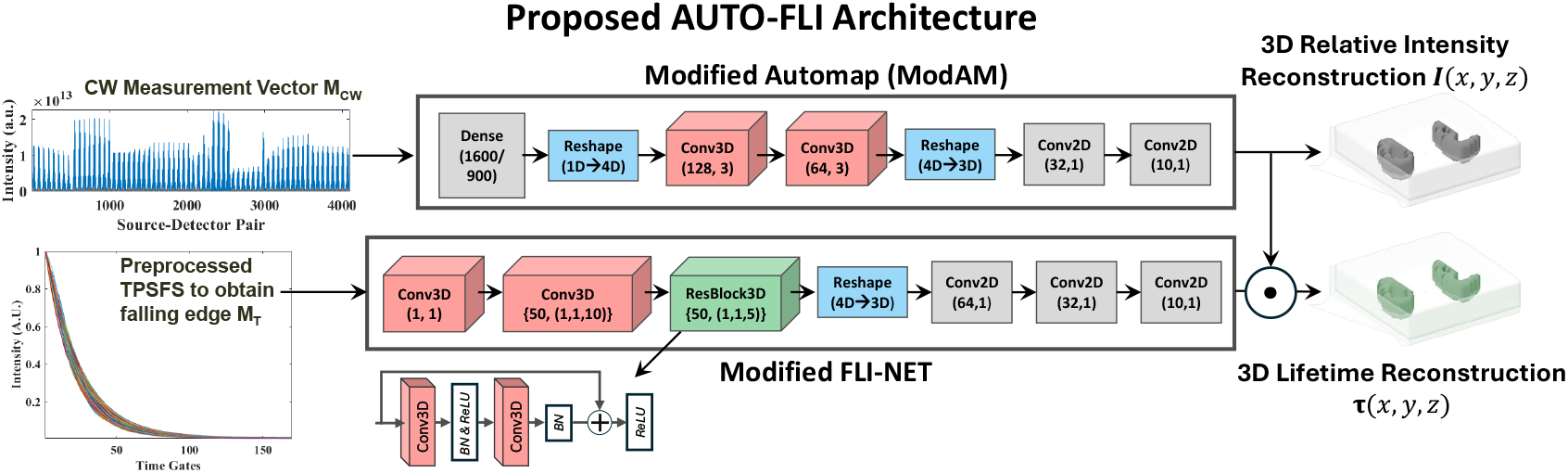
The Proposed AUTO-FLI architecture, a combination of ModAM (top) and modified FLI-NET (bottom).

The ModAM branch begins with a Dense layer 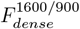, which is particularly useful for mapping the one-dimensional CW measurement vector *M*_*CW*_ ∈ 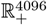 to a lower-dimensional feature representation. This layer helps capture the complex relationship between the measured CW signal and the underlying spatial distribution of fluorescent inclusions, facilitating the reconstruction of their three-dimensional morphology in subsequent stages of the network. Furthermore, the Dense layer, with ReLU activation, serves as a projection transformation, whose number of hidden neurons depends on the specific size of the observed object; that is, it depends on the Field-of-View (FOV). The FOV for the phantom is 40 × 40 mm^2^, and for the mouse is 30 × 30 mm^2^, resulting in 1600 and 900 neurons, respectively. Next, the resulting feature tensor is reshaped into a 4D tensor to prepare it for 3D convolutional (Conv3D) operations:

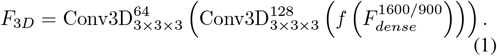

Here, and in all subsequent equations, *f* (·) indicates that after reshaping, an implicit function of the argument tensor (including batch normalization and ReLU) is applied before passing it on to the next layer. The output of the Conv3D layers *F*_3*D*_ is then reshaped back to a 3D volume and refined by 2D convolutional (Conv2D) layers:

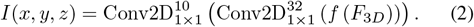

The final output of this branch is the 3D intensity volume *I*(*x, y, z*), which indicates the spatial distribution of the fluorescence intensity (quantum yield) within the volume of interest.

The Modified FLI-NET branch extracts spatiotemporal features from preprocessed, normalized time-resolved signals and reconstructs a 3D lifetime volume. Specifically, it operates on the falling edges of the TPSFs measured for each illumination pattern and each pixel on the tissue or phantom surface. This preprocessed input is denoted by 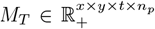 are the number of illumination patterns. First, a sum aggregation *F*_*Agg*_, with a learnable weighting factor *α*, is applied to the illumination patterns:

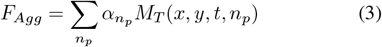

This step is mathematically equivalent to a 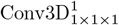 operation.

Next, a Conv3D layer with a kernel size of 1 × 1 × 10 (where *k* is the kernel size along the temporal axis) is deployed to capture spatially independent features along the TPSFs:

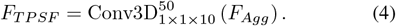

A 3D Residual Block (ResBlock3D) further enriches temporal feature extraction without substantially increasing parameter count compared to a standard deep convolutional network stacking multiple Conv3D layers sequentially [30]. Mathematically, the ResBlock3D includes two internal Conv3D layers:

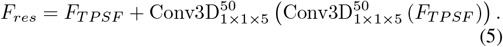

The output *F*_*res*_ is reshaped into a 3D volume and passed through a series of Conv2D layers to produce an unmasked 3D lifetime map *τ*_*raw*_:

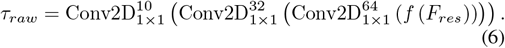

Finally, we constrain the raw lifetime map *τ*_*raw*_(*x, y, z*) by the 3D intensity volume *I*(*x, y, z*), ensuring lifetime values are only assigned to regions of fluorescent activity:

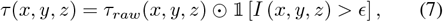

where 𝕝 is an indicator function, ⨀ is element-wise multiplication, and *ϵ* = 0.5 is a standard threshold for normalized intensity data in the field of optical tomography [26].

Figure 2 shows the network architecture and the parameters of the different network layers used for training on the experimental tissue-mimicking phantom and the mouse phantom. Furthermore, Table I provides details of network training. We trained an ensemble of 30 independent networks on a dataset of 1, 500 samples, divided into training, validation, and test subsets following an 80*/*15*/*5 split. All networks were optimized using the Adam optimizer, employing mean squared error (MSE) loss computed separately for the relative intensity and lifetime estimates against their ground-truth values. We varied the relative concentration, depth, thickness, and lifetime of the *in silico* phantoms in ranges relevant to soft tissue, ensuring robust performance under a variety of realistic conditions.

**TABLE 1.**
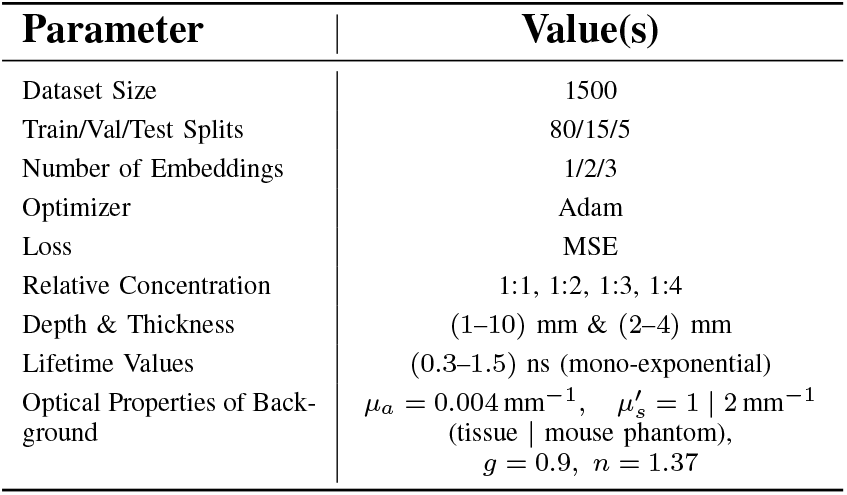
NETWORK TRAINING DETAILS (GPU: GEFORCE RTX 2080 TI)

### C. Experimental Setup

The experimental setup consists of a gated-ICCD system. The optical imaging is performed in reflective geometry with a FOV of approximately 40 × 40 mm^2^. A total of 201 gates are used with a gatewidth of 40 ps. The *k*-space illumination patterns are projected using a Digital Micromirror Device (D4110, Digital Light Innovations, TX). In the case of the mouse phantom, the FOV is adjusted to approximately 30 ×30 mm^2^ to project the patterns onto a flat, central area on the mouse’s abdomen. For imaging the experimental phantoms, we excite at a wavelength of 700 nm using a Mai Tai high-powered laser (Spectra-Physics, CA). Further details of the experimental setup can be found elsewhere [8].

### D. Phantom Preparation

To prepare the tissue-mimicking phantom, we combine distilled water, 1% India Ink (Speedball Art Products, NC), 20% intralipid (Sigma–Aldrich, MO) of volumes of 157.05 ml, ml, and 11.90 ml, respectively, with 1.7 g of agar to form a homogeneous phantom that has roughly the same background OPs as the *in silico* phantoms used in training. Two cylindrical cavities are formed in the phantom at depths of (2–5) mm, with the centers approximately 20 mm apart, in which we contain AF700 at 50 *μ*M concentration in Dimethyl sulfoxide (DMSO) and Phosphate-buffered saline (PBS) buffers.

Additionally, we have previously developed and optimized tissue-mimicking phantoms with controlled 3D fluorescent inclusions [31]. For this study, we fabricated a mouse-shaped tissue-mimicking phantom by casting a postmortem mouse in Plaster of Paris and creating durable polydimethylsiloxane (PDMS) phantoms with anatomically accurate surface features. The OPs were tuned using titanium dioxide and India ink to achieve physiologically relevant scattering and absorption characteristics. Next, we embedded three small glass capillaries (inner radius: 2 mm; outer radius: 2.5 mm) filled with AF700 at a fixed concentration of 10 *μ*M. Two capillaries (one in PBS and one in DMSO) were positioned at a shallow depth (1–2) mm near the mammary fat pad region, while the third capillary (AF700 in PBS) was placed at a deeper location (3–4) mm in the liver region. The overall dimensions of the mouse phantom were approximately 3×3×9 cm^3^. The background OPs of the phantom were measured using Spatial Frequency Domain Imaging and determined to be 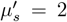 mm^*−*1^ and *μ*_*a*_ = 0.004 mm^*−*1^. During training, these estimated OPs were incorporated into the DL network to improve model accuracy.

## III. Results

### A. In Silico Results

The performance of our proposed AUTO-FLI network is first tested on *in silico* phantoms that were not part of the training stage. For demonstration, we used an *in silico* phantom containing two fluorescent inclusions, each of which has a different lifetime (0.9 ns for the left inclusion and 0.7 ns for the right inclusion) and different relative concentration (1 : 2). The inclusions are placed at depths of (2–4) mm, as shown in Fig. 3A. The 3D lifetime reconstruction by the AUTO-FLI network, rendered by the Imaris software, is displayed in Fig. 3B. In Fig. 3C, we display the Ground Truth (GT) relative concentration map, and in Fig. 3D, we show the obtained 2D intensity maps corresponding to depths (1–4) mm. Additionally, in Fig. 3E, we present the intensity profile along the red dotted line in Fig. 3C. It is evident that the ModAM branch of the AUTO-FLI network reconstructs the 3D intensity with a high degree of fidelity and manages to recover the relative concentration accurately. The accuracy is numerically validated by a low VE of 17.86% and an MSE of 0.0328. The 3D lifetime reconstruction results are shown in Figs. 3F-H. The 3D reconstruction, in terms of structure, is the same as the output of the ModAM branch since the output of the 3D FLI-NET branch is constrained by the 3D intensity mask. In Fig. 3G, we display the 2D lifetime maps corresponding to depths (1–4) mm. The GT lifetime map is illustrated for comparison in Fig. 3F. Additionally, we present the violin plot distribution of the reconstructed lifetime values for the two fluorescent species (along with the GT) in Fig. 3H. It can be deduced from these plots that the proposed AUTO-FLI framework accurately reconstructs the 3D lifetime (with an MSE value of 0.0088).

**Fig. 3.**
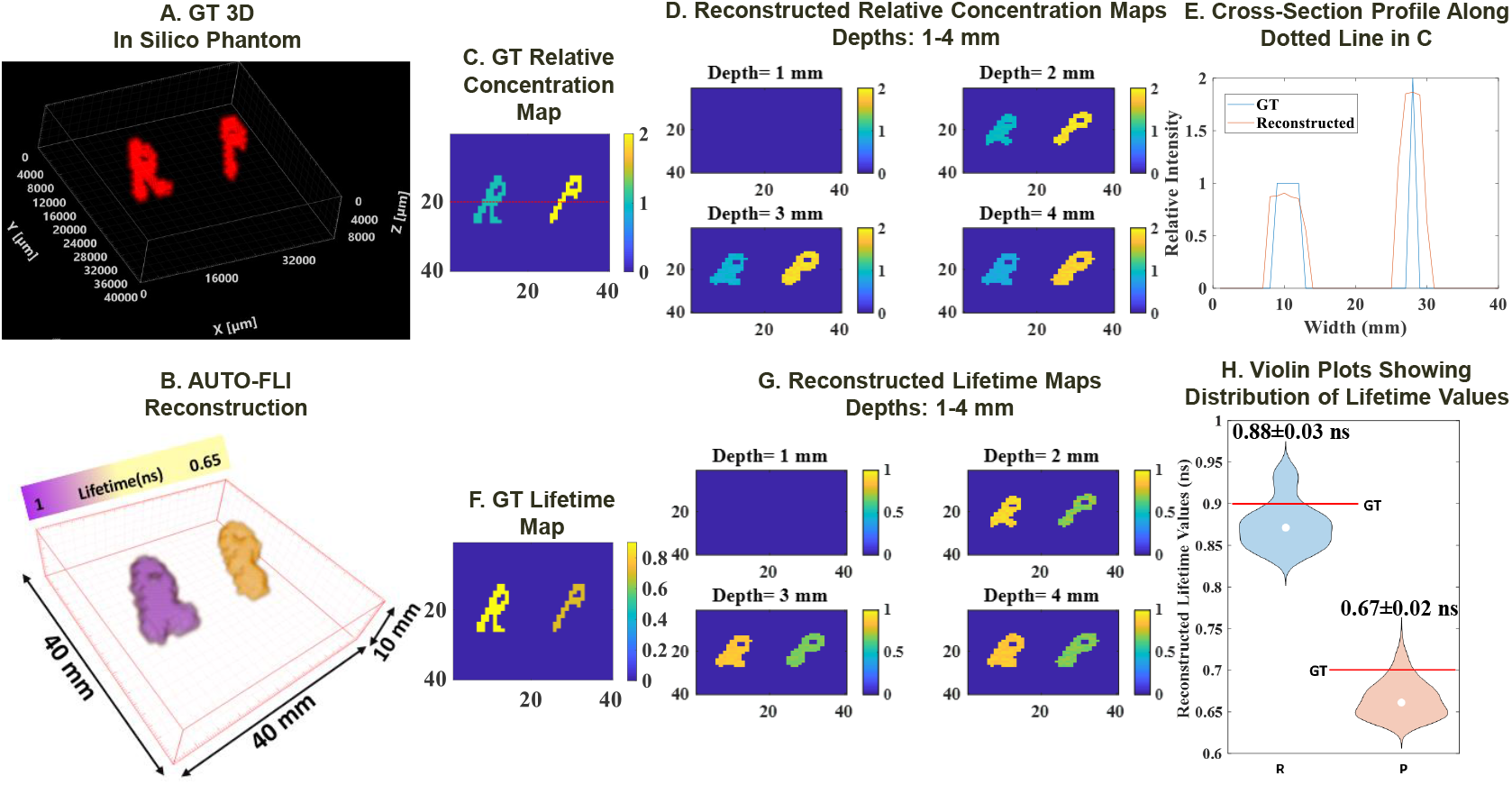
The GT 3D *in silico* phantom, not part of the training stage of the AUTO-FLI network, is shown in A with the GT intensity and lifetime maps in C and E, respectively. The 3D reconstruction, rendered by Imaris, by the AUTO-FLI framework is displayed in B, and the reconstructed 2D intensity maps and 2D lifetime maps are in D and F, respectively. The intensity profile along the red dotted line along with the GT is plotted in E, while the violin plots showing the reconstructed lifetime distribution of the two fluorescent species (with red lines representing the GT) are presented in H. The 3D reconstruction results are shown at **50%** of the maximum isovolume.

### B. Experimental Phantom Results

To further validate the proposed AUTO-FLI network, we tested its performance on an experimental tissue-mimicking phantom. The same network trained for the *in silico* phantoms is used to reconstruct the experimental phantom. A schematic of the phantom is shown in Fig. 4A. The experimental phantom consists of two cylindrical fluorescent inclusions AF700 in DMSO (left) and PBS (right) embedded in agar (background OPs set using a mixture of ink and intralipid), the center of which is placed at a depth of 3 mm from the surface of the phantom. The inclusions are approximately 4 mm thick. A schematic of the phantom is shown in Fig. 4A. The two different buffers ensure that the two inclusions have slightly different lifetimes while having approximately the same concentration. However, it should be noted that DMSO gives a slightly brighter signal than PBS. The results of the 3D volumetric reconstruction, again rendered by Imaris, are presented in Fig. 4B, and the 2D normalized intensity maps corresponding to depths (1–6) mm are illustrated in Fig. 4C. Moreover, the violin plots showing the distribution of the relative intensity values are plotted in Fig. 4D. It is observed that the ModAM branch of the network reconstructs the relative concentration close to what is expected, with a slightly higher concentration of AF700 in DMSO (left) compared to AF700 in PBS (right). The relative concentration (average) is obtained as 1.08 : 1 (DMSO:PBS). The 2D lifetime maps reconstructed (corresponding to depth (1–6) mm) by the AUTO-FLI framework are shown in Fig. 4E, while the distribution of lifetime values are presented in Fig. 4F. The reconstructed lifetime values agree closely with the lifetime values for AF700 in DMSO and PBS quoted in the literature [32]. Furthermore, for benchmarking, we obtain the lifetime values estimated from the experimental phantom using the state-of-the-art AlliGator software. The AlliGator software estimates the lifetime of a 2D mask placed on the 2D widefield image of the phantom. The mean and standard deviation of the left (DMSO) and right (PBS) inclusion is calculated by AlliGator as (1.14±0.02) ns and (1.03±0.02) ns, respectively. Thus, the lifetime values obtained from AlliGator are comparable to those produced by AUTO-FLI. However, it is to be noted that the AUTO-FLI network can carry out a single 3D lifetime reconstruction in approximately 1 s (NVIDIA GeForce RTX 2080 Ti). Conversely, on the same GPU, it takes almost 10 minutes for AlliGator to calculate the lifetime on 10 pixels. Hence, the results justify the suitability of the proposed workflow for experimental phantoms.

**Fig. 4.**
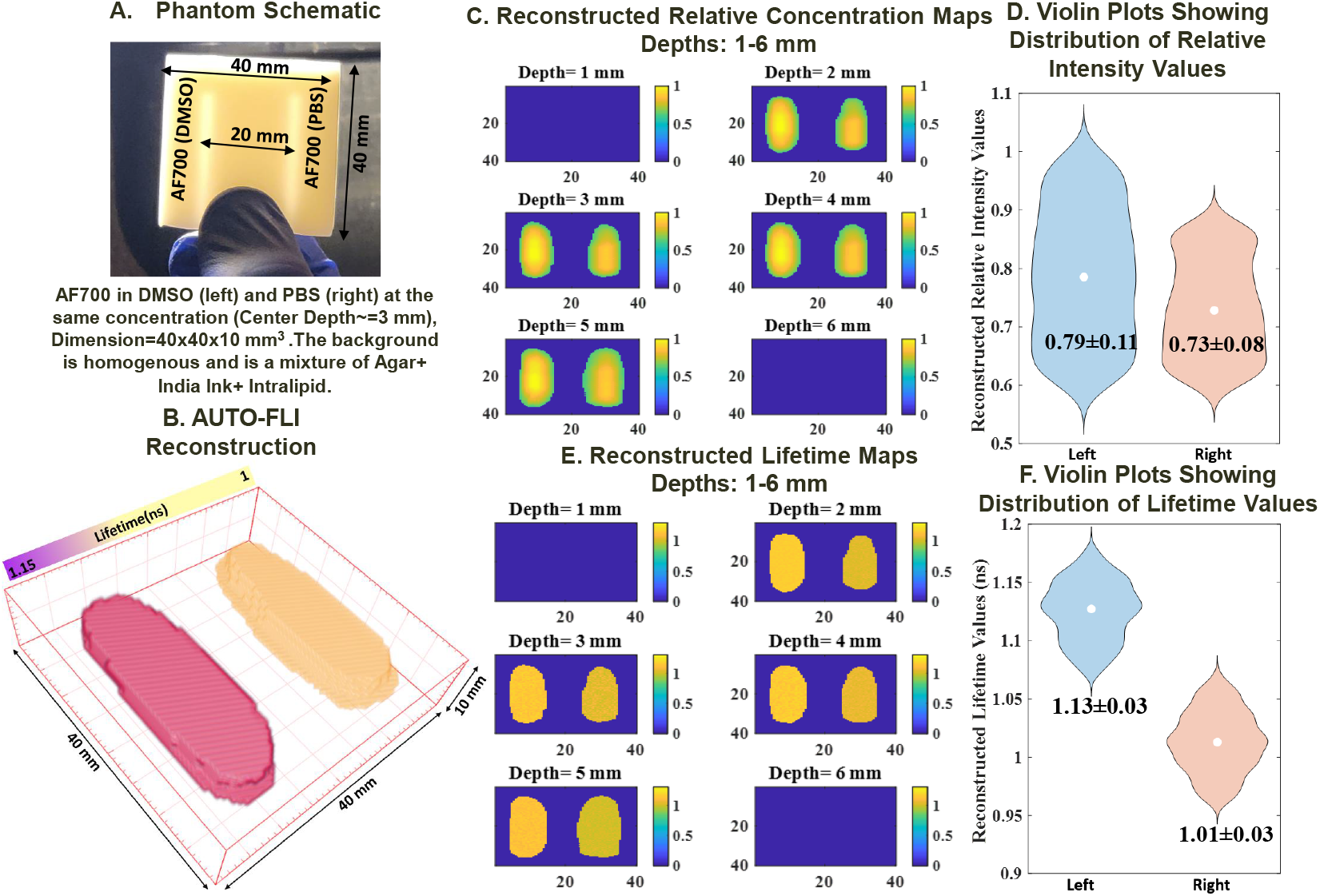
The schematic of the experimental phantom is shown in A. The 3D reconstruction (rendered by Imaris) is presented in B, along with the obtained 2D relative concentration and lifetime maps in C and E, respectively. The distribution of the relative intensity and lifetime values, along with the mean and the standard deviation, for both fluorescent inclusions are presented in terms of violin plots in D and F, respectively. The 3D reconstruction results are shown at **50%** of the maximum isovolume.

### C. Mouse Phantom Results

To demonstrate the strength of our algorithm and its potential translatability to pre-clinical scenarios, such as in small animal imaging, we present the results on a biologically accurate mouse-mimicking phantom. As mentioned before, the data generator and the network are suitably adjusted to account for the mouse phantom’s different physical dimensions and OPs. In Fig. 5A, we see the 3D reconstruction results of the lifetime generated by AUTO-FLI rendered on Imaris. As expected, the network resolves the structure of the two shallow tubes very well and, to a slightly lesser extent, the deeper tube. Figs. 5B and 5C show the average 2D intensity and lifetime maps generated by the two branches of the AUTO-FLI network. These results are demonstrated graphically in the form of violin plots in Figs. 5D and 5E. The results agree with the expectation of slightly higher intensity for the DMSO tube, similar intensities for the two PBS tubes, and a higher lifetime for the DMSO tube. The lifetime results are benchmarked against AlliGator (average of 1.10 ns and 1.00 ns for tubes 1 and 3, respectively, and 1.80 ns for tube 3) and found to be in good agreement. It is again to be noted that AlliGator only produces results on a 2D mask, and the computational time is manifold higher. These results show that this network may be utilized in pre-clinical applications.

**Fig. 5.**
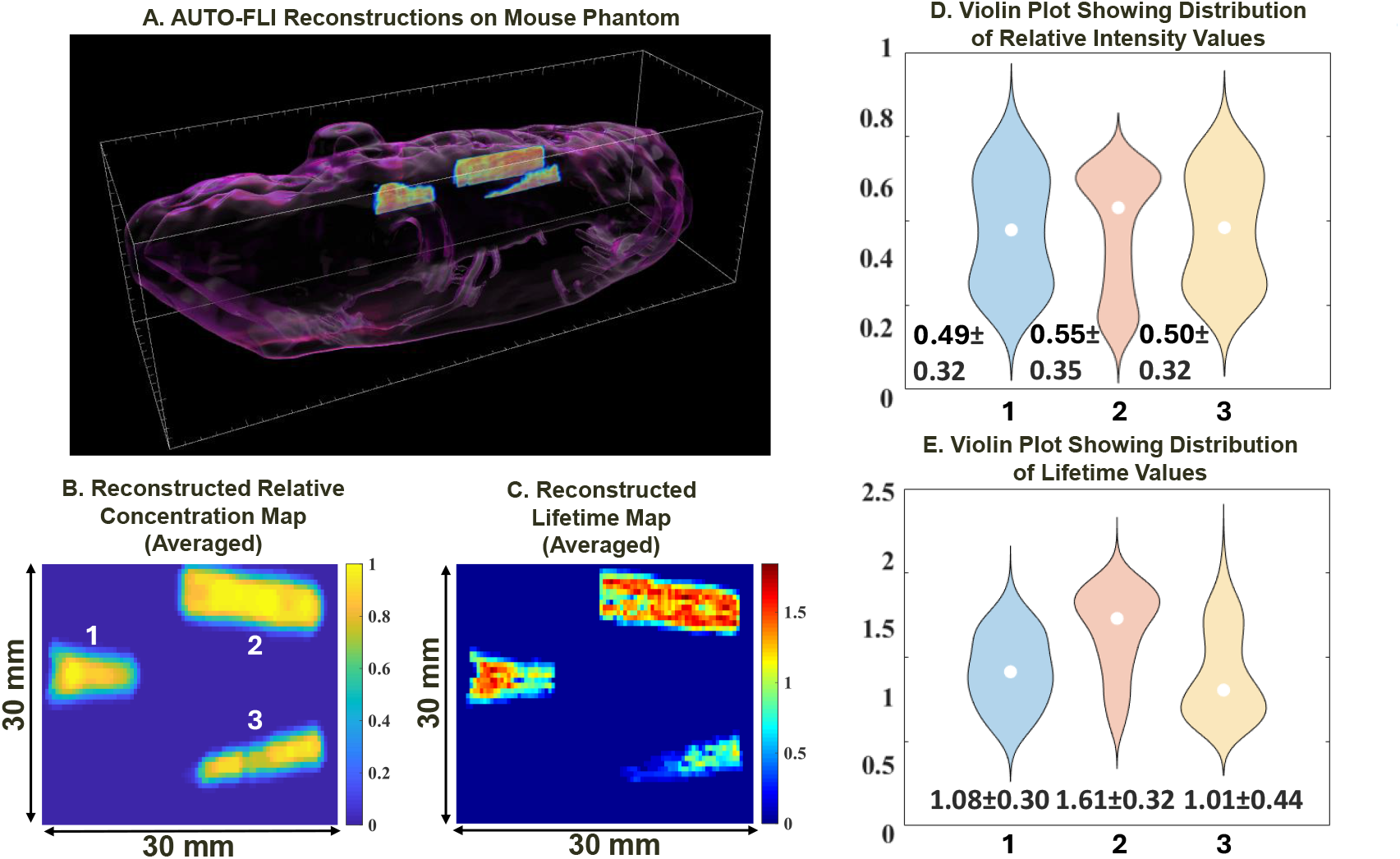
The 3D reconstruction (rendered by Imaris) is presented in A, along with the obtained average 2D relative concentration and lifetime maps in B and C, respectively. The distribution of the relative intensity and lifetime values, along with the mean and the standard deviation, for both fluorescent inclusions are presented in terms of violin plots in D and E, respectively. The 3D reconstruction results are shown at **50%** of the maximum isovolume.

## IV. Discussion

This work presents a novel and comprehensive framework—AUTO-FLI—for simultaneous 3D intensity and fluorescence lifetime reconstructions using a two-stage DNN architecture. The design of the AUTO-FLI system represents a significant step forward in FLT, addressing long-standing challenges such as the dual ill-posed nature of the inverse problem, high computational demands, and limited scalability of classical methods.

One of the key innovations lies in the decoupling of intensity and lifetime estimation via the two-stage approach. This design leverages the strength of modular learning by first accurately localizing fluorescent regions through the ModAM branch and subsequently constraining lifetime estimation using the modified FLI-NET. Our results demonstrate that this separation improves stability and reconstruction fidelity and mitigates noise propagation between spatial and temporal domains. This finding aligns with the concept of multi-task learning [33], which posits that breaking complex problems into simpler, focused subtasks improves overall model performance.

In our case, this is supported by the ablation experiment shown in Fig. 6, where removing the ModAM branch and relying solely on the FLI-NET branch leads to degraded reconstructions in both intensity and lifetime domains. Although the network can still capture the basic structure, the lack of spatial priors results in higher reconstruction errors, underscoring the critical role of intensity-guided lifetime estimation. The superior performance of the AUTO-FLI approach was evidenced by markedly lower MSE and volume error (VE) metrics compared to the single-stage alternative (VE: 17.86% vs. 44.26%, MSE: 0.0088 vs. 0.023), validating the importance of staged reconstruction.

**Fig. 6.**
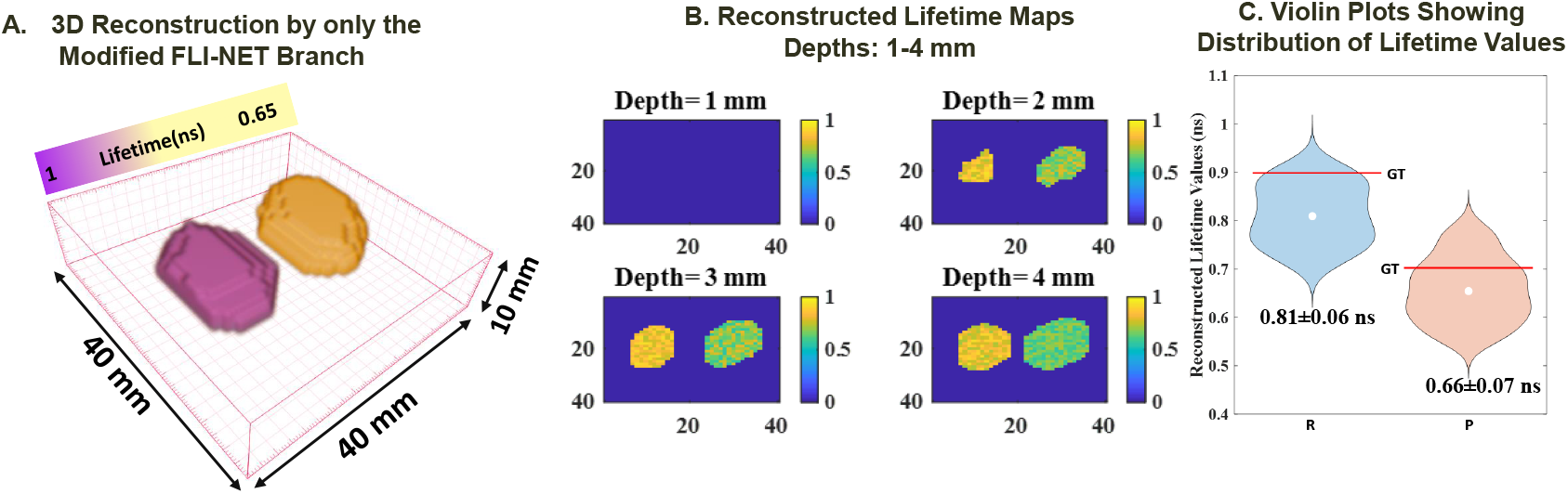
The 3D reconstruction obtained using only the Modified FLI-NET stage (by eliminating ModAM) is shown in A for the same *in silico* phantom used in Fig. 3 along with the obtained 2D lifetime maps in B, and the violin plot distribution of the reconstructed lifetime values in C. Again, the 3D reconstruction results are shown at **50%** of the maximum isovolume.

Additionally, AUTO-FLI offers substantial advantages in computational efficiency. A full 3D lifetime reconstruction is achieved in under one second on an RTX 2080 Ti GPU, orders of magnitude faster than classical tools like AlliGator, which require several minutes for just a 2D region of interest. This performance enables real-time or near-real-time applications, making AUTO-FLI particularly suited for time-sensitive biomedical workflows such as fluorescence-guided surgery.

The generalizability of the system has been established in three validation levels: *in silico* data, a tissue-mimicking phantom, and a biologically realistic small animal phantom. The method agreed with ground truth values and demonstrated robustness under varying fluorophore concentrations, depths, and tissue OPs. Unlike prior deep learning-based FLT methods often relying on oversimplified geometries or 2D assumptions, AUTO-FLI successfully resolves complex 3D structures with sub-nanosecond lifetime precision, validated against the benchmark AlliGator software.

To assess the consistency of the method, we conducted a stability study by training AUTO-FLI 30 times using randomized initializations. The training and validation curves, shown in Fig. 7, exhibit smooth convergence with a low standard deviation (±0.005), demonstrating the robustness and repeatability of the proposed architecture.

**Fig. 7.**
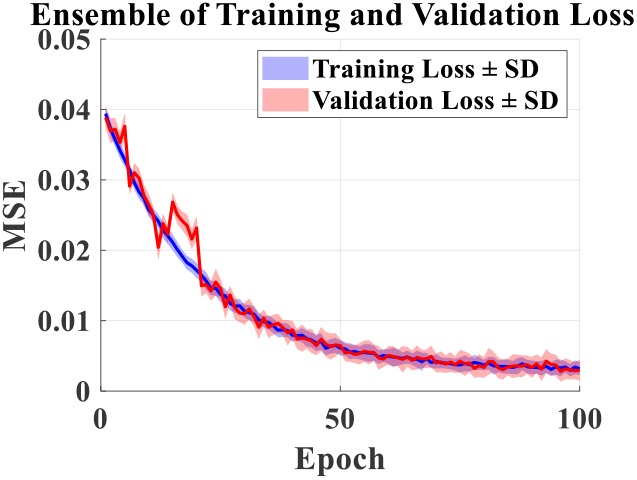
An ensemble of training and validation curves obtained by training the AUTO-FLI network **30** times.

The *in silico* data generation workflow, driven by MCX-based photon propagation modeling, further reinforces the method’s fidelity. By incorporating accurate physics-based simulation of scattering, absorption, and experimentally measured IRFs, the network benefits from training on data that closely reflects experimental conditions. This approach circumvents the need for large experimental datasets and enhances reproducibility—two major limitations in prior deep learning-based optical imaging workflows.

Compared to traditional approaches that use asymptotic approximations, CT-based structural priors, or linearized inverse models, AUTO-FLI provides a self-contained, data-driven solution that offers superior reconstruction quality and speed without sacrificing physical realism. Although recent efforts using physics-informed neural networks (PINNs) have shown promise [34], they often require extensive computational resources and expert domain knowledge to integrate governing equations. AUTO-FLI instead combines a physics-informed data generation backend with a high-performing, conventional CNN, achieving an optimal trade-off between realism, generalizability, and accessibility.

Despite its rigorous validation, the proposed work has some limitations. We used a simplified mono-exponential model for lifetime estimation. In contrast, the original FLI-NET model employed a bi-exponential model that was better suited for various applications, including multiplexed and FRET studies. Future research will focus on extending the workflow to accommodate such higher-order exponential models. Furthermore, this study concentrated on establishing and validating the novel 3D FLI technique in well-controlled environments, using accurate synthetic datasets that cover a broad parameter space and an anatomically precise small animal phantom. Subsequent studies will aim to apply this technique in *in vivo* settings. These applications present additional challenges, such as potential changes in the optical properties of tissues, variability in tissue boundaries, and the need for robust validation methods. Controlling lifetime contrasts spatially in 3D is particularly challenging in biological systems, making it difficult to establish a reliable ground truth. Therefore, lifetime quantification validation should involve tissue extraction and *ex vivo* validation, carefully considering the microenvironmental parameters that affect lifetime values. Such a workflow will require a comprehensive study beyond the current scope of this work.

## V. Conclusion

In this study, we introduced AUTO-FLI, an innovative deep learning (DL) framework for 3D fluorescence lifetime tomography (FLT) that significantly enhances accuracy and computational efficiency. Using a two-stage convolutional neural network (CNN) architecture, AUTO-FLI enables high-fidelity reconstructions of 3D intensity and lifetime distributions from raw 2D fluorescence decay data. A key feature of our approach is integrating a versatile data generator that leverages an accurate light propagation model in highly scattering media, reducing training times and resource consumption while improving the model’s robustness. Overall, AUTO-FLI demonstrates promising potential for advancing preclinical research and clinical diagnostics.

## Acknowledgment

The authors would like to thank Mr. Harrison Yee for his help with Imaris.

## Notes

This work was supported by the funding received from the National Institutes of Health under grants R01- CA271371, R01-CA237267, and R01-CA250636.

### Competing Interest Statement

The authors have declared no competing interest.

### Summary of Updates

Methods Section has been updated to include the mathematical framework for AUTO-FLI. Results Section has been updated to include mouse-mimicking phantom

